# Ketamine-induced NMDA receptor hypofunction alters social and locomotor behavior in adult zebrafish

**DOI:** 10.1101/2025.09.16.676572

**Authors:** Matheus Gallas-Lopes, Daniela V. Müller, Thailana Stahlhofer-Buss, Leonardo M. Bastos, Sofia Z. Becker, Samara M. Bruck, Angelo Piato, Ana P. Herrmann

**Affiliations:** Laboratório de Neurobiologia e Psicofarmacologia Experimental (PsychoLab), Departamento de Farmacologia, Instituto de Ciências Básicas da Saúde (ICBS), Universidade Federal do Rio Grande do Sul (UFRGS), Rua Ramiro Barcelos 2600/430, Porto Alegre, Rio Grande do Sul, 90610-264, Brazil; Laboratório de Psicofarmacologia e Comportamento (LAPCOM), Departamento de Farmacologia, Instituto de Ciências Básicas da Saúde (ICBS), Universidade Federal do Rio Grande do Sul (UFRGS), Rua Ramiro Barcelos 2600/411, Porto Alegre, RS, 90610-264, Brazil; Programa de Pós-Graduação em Farmacologia e Terapêutica, Departamento de Farmacologia, Instituto de Ciências Básicas da Saúde (ICBS), Universidade Federal do Rio Grande do Sul (UFRGS), Rua Ramiro Barcelos 2600/625, Porto Alegre, RS, 90610-264, Brazil

**Keywords:** Ketamine, NMDA receptor hypofunction, Social withdrawal, Hyperlocomotion, Stereotyped swimming, Schizophrenia-like phenotypes

## Abstract

**Background:** NMDA receptor antagonists, such as ketamine, are widely used to model schizophrenia-related phenotypes in preclinical studies. While zebrafish have emerged as a promising model organism for neuropsychiatric research, few studies have characterized their behavioral responses to repeated ketamine exposure.

**Methods:** Three independent experiments were conducted to evaluate the acute, repeated and sustained behavioral effects of ketamine in adult zebrafish. In Experiment I, fish were exposed to 10, 20, or 40 mg/L ketamine once daily for five days, and submitted to the social preference (SPT) and open tank (OTT) tests on days 1 and 5, and re-exposed and re-tested on day 7 after a 48-hour washout. Experiments II and III assessed whether behavioral changes persisted following 5- or 14-day exposure protocols, with testing conducted 48 hours after the final treatment.

**Results:** Ketamine induced robust, concentration-dependent alterations in Experiment I: it reduced social interaction and increased locomotor activity in the SPT on all experimental days, while increased rotational behavior in the OTT on days 1 and 5. These effects did not intensify over repeated exposure and were not sustained after a 48-hour washout in either protocol (Experiments II and III).

**Conclusions:** The results support the utility of zebrafish for modeling acute behavioral responses to NMDA receptor antagonism, capturing features of schizophrenia-like phenotypes. However, no evidence of behavioral sensitization or lasting disruption was observed, diverging from rodent studies. Future studies should incorporate antipsychotic validation, neurochemical analyses, and alternative exposure strategies to further develop zebrafish as a translational model for psychiatric research.

## INTRODUCTION

Animal models that recapitulate schizophrenia-like behavioral phenotypes are essential for uncovering the disorder’s mechanisms and advancing translational research in psychiatry. Among the pharmacological tools used in preclinical studies, non-competitive N-methyl-D-aspartate receptor (NMDAR) antagonists such as MK-801, phencyclidine, and ketamine are widely employed to induce behavioral alterations that reflect core symptom domains of schizophrenia [1–5]. By mimicking NMDAR hypofunction, these compounds contribute to dopaminergic hyperactivity in subcortical regions and evoke transient behavioral changes that resemble the positive, negative, and cognitive symptoms associated with the disorder [2, 5–8].

Rodent models have played a central role in preclinical schizophrenia research. However, a substantial number of pharmacological candidates that showed promise in these models have failed in clinical trials, highlighting a critical gap in translational success [9, 10]. These failures raise concerns about the external validity of such models, particularly their ability to predict complex human outcomes. In response, there is growing recognition of the need for alternative models that can enhance translational relevance and reduce species-specific biases. One such strategy involves testing conserved behavioral and pharmacological effects across diverse species [11–14]. Among non-rodent models, zebrafish (*Danio rerio* Hamilton, 1882) have emerged as a valuable organism in neurobehavioral research due to their well-characterized nervous system, evolutionary conservation of key neurotransmitter pathways, and suitability for high-throughput experimental designs [15–17]. These features make zebrafish a promising platform for cross-species validation and for improving the predictive value of preclinical findings.

Previous work from our group demonstrated that the NMDAR antagonist MK-801 (dizocilpine) disrupts social behavior and induces hyperlocomotion in adult zebrafish, supporting the model’s relevance for studying schizophrenia-like phenotypes [18, 19]. Compared to MK-801, the behavioral effects of ketamine in zebrafish are less consistently described. Although ketamine is widely used in preclinical models of schizophrenia, studies in zebrafish vary considerably in design and outcomes tested, with limited agreement on its impact across social, locomotor, and exploratory domains [1].

The effects of ketamine exposure on zebrafish behavior have been most commonly examined after acute exposures [1, 20–22], and far less is known about how behavior changes with repeated administration, or about whether these effects persist once treatment has ended. In rodents, repeated exposure to NMDAR antagonists like ketamine results in progressively heightened behavioral responses or long-lasting changes that remain after drug clearance [23–28]. These findings suggest that both the temporal dynamics and the cumulative impact of ketamine exposure are important factors in shaping behavioral outcomes. Whether zebrafish show similar patterns is still unknown.

This study aimed to characterize schizophrenia-like behavioral effects of ketamine in adult zebrafish across three experimental protocols. Experiment I examined how behavioral responses evolved over repeated exposures and test sessions to assess the potential development of sensitization. Experiment II evaluated whether ketamine-induced behavioral alterations persisted after a short-term (5-day) exposure followed by a 48-hour washout period. Experiment III extended this investigation by assessing whether a longer-term (14-day) exposure regimen would lead to detectable effects after the same washout interval. Together, these protocols allowed for the systematic assessment of both short- and long-term consequences of ketamine exposure across different behavioral domains.

## MATERIALS AND METHODS

This study was conducted in accordance with ethical guidelines and approved by the institutional animal ethics committee of the Universidade Federal do Rio Grande do Sul (approval: #35525/2018). The reporting of this study follows the ARRIVE guidelines for animal research transparency and rigor [29]. A completed ARRIVE Essential 10 checklist is available at osf.io/yqv6a. The experimental protocols for all three experiments were preregistered on the Open Science Framework, specifying hypotheses, dependent variables, conditions, analyses, outlier exclusion criteria, and sample size calculation [30] (Experiment I: osf.io/r7xdy [31]; Experiment II: osf.io/kjf3r [32]; Experiment III: osf.io/vkcft [33]). Assistance from ChatGPT-4o (OpenAI, San Francisco, CA, USA) was used to support the optimization of text and formatting throughout the manuscript. All outputs were critically reviewed, edited, and validated by the authors, who take full responsibility for the content and interpretation of the work.

### Animals, housing, and husbandry

A total of 224 adult wild-type short-fin zebrafish, aged between 3 and 4 months, were used across three separate experiments: 128 animals in Experiment I, and 48 animals each in Experiments II and III. Of these, 217 completed the full experimental protocol and behavioral assessments (Experiment I: 122; Experiment II: 48; Experiment III: 47) – missing data is due to the death of 7 fish during the course of the experiments (full details in the “Statistical analysis” section). The final sample included 111 females and 106 males, with the following sex distribution: Experiment I – 62 females, 60 males; Experiment II – 25 females, 23 males; Experiment III – 24 females, 23 males. As no sex-specific effects were found in Experiment I, data from male and female animals were combined in all analyses. Animals were sourced from a commercial supplier (Delphis, Porto Alegre, RS, Brazil). All animals were maintained at the zebrafish facility of the Instituto de Ciências Básicas da Saúde (Universidade Federal do Rio Grande do Sul) under a controlled 14:10 hour light/dark cycle (lights on at 07:00). Prior to the onset of experiments, fish were allowed to acclimate to laboratory conditions for at least two weeks. They were group-housed in 16 L glass tanks (40 × 20 × 24 cm) enriched with small PVC tubes and kept at a maximum density of two animals per liter. Tanks were filled with dechlorinated tap water and equipped with continuous aeration and mechanical, biological, and chemical filtration systems (Altamar, Jacareí, SP, Brazil). Water quality parameters were maintained within acceptable ranges for zebrafish: temperature at 27 ± 2 °C, pH 6.8 ± 0.3, and conductivity between 500–800 μS/cm. Fish were fed twice daily with a combination of *Artemia salina* and commercial flake food (Poytara, Araraquara, SP, Brazil). During the experimental phase, animals were redistributed into smaller aquariums (18 × 18 × 18 cm) according to the treatment group. Each group was housed separately, and all tanks were kept under identical conditions within the same facility, including lighting, temperature, and vertical level. Tanks were coded to maintain caregiver blinding regarding treatment allocation. There were two housing tanks for each treatment group in all experiments. Physical barriers were used to prevent visual contact between groups, both in housing aquariums and during exposure to treatments. At the conclusion of the experiments, animals were euthanized using hypothermic shock. This procedure involved immersion in water cooled to 2–4 °C for at least two minutes after cessation of opercular movements, followed by decapitation to ensure death. Sex was confirmed post-mortem via examination of the gonads.

### Drugs

Ketamine solutions were prepared by diluting a 10% (v/v) injectable formulation (Syntec, Tamboré, SP, Brazil) in water maintained under the same conditions as the housing tanks. In Experiment I, concentrations of 10, 20, and 40 mg/L were used. In Experiments II and III, only the 40 mg/L concentration was used. These concentrations were selected based on previous studies reported in the literature [1, 21, 22]. Fish were exposed to the treatment solutions by immersion in a beaker for 20 minutes. The 20-minute immersion period was chosen following established protocols reported in the literature and pilot experiments conducted by our group using another NMDA receptor antagonist in zebrafish [1, 18, 21, 22]. Control animals were handled in the same way but immersed in ketamine-free water under identical conditions.

### Experimental Design

This study included three independent experiments, each conducted with a separate set of animals. The design and timeline for each experiment are illustrated in Figure 1. Animals were allocated to the experimental groups using block randomization, ensuring balance across presumed sex and home tank origin. Because zebrafish sex cannot be confirmed without gonadal inspection, group allocation was counterbalanced based on external sex characteristics at the time of assignment. To minimize potential biases, the order of behavioral testing was determined using block randomization, with the sequence of randomized blocks generated via random.org, taking into account the test apparatuses to ensure even distribution across treatment conditions. The experimental unit in all analyses was a single animal, with each fish contributing one independent data point to the statistical comparisons. Both the experimenters conducting the behavioral assessments and those responsible for transferring the animals to the test apparatus were blinded to group allocation.

**Figure 1.**
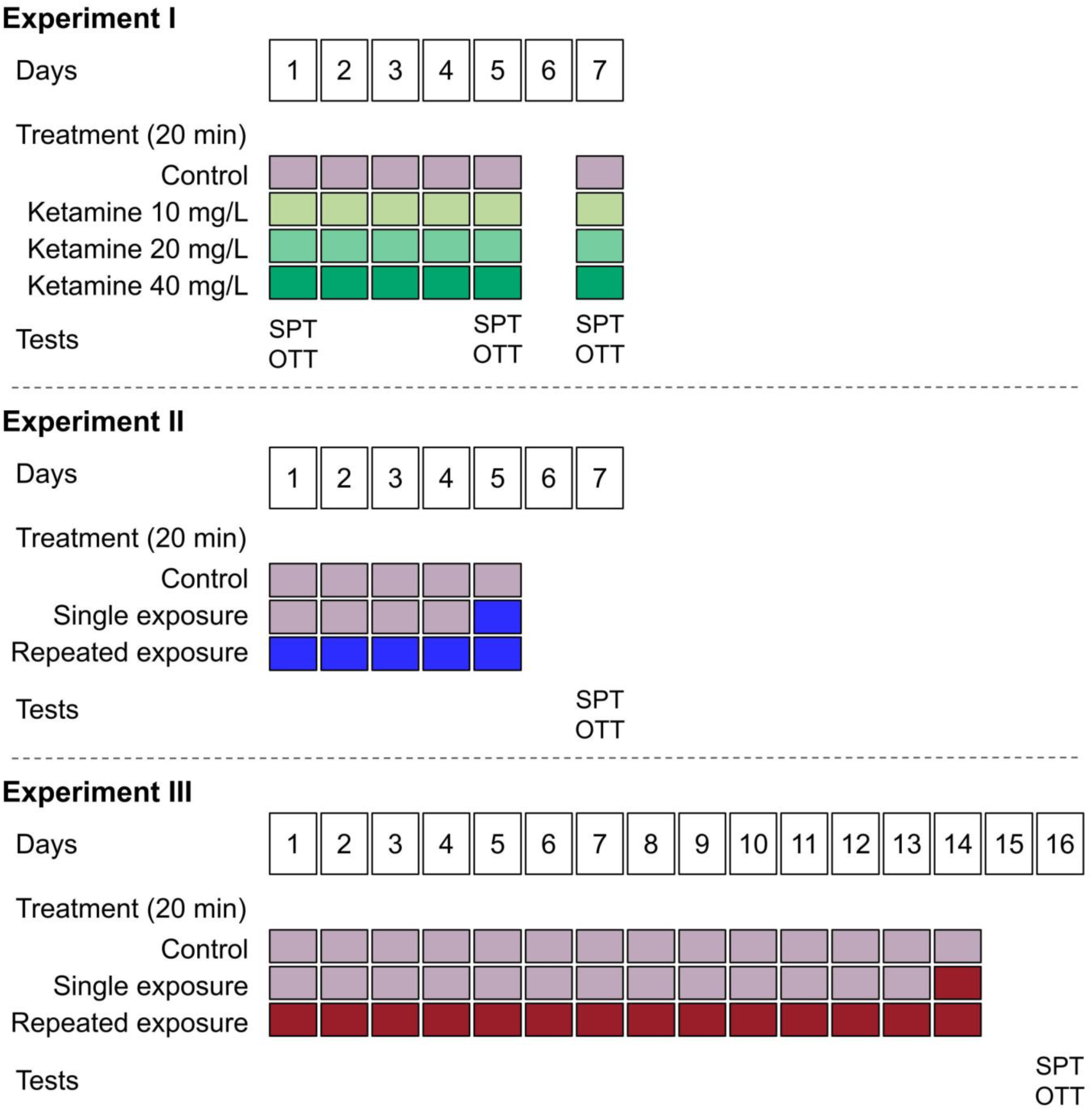
Overview of the experimental design evaluating the behavioral effects of ketamine exposure in zebrafish. Experiment I: fish were exposed via immersion to water or ketamine (10, 20, or 40 mg/L) for 20 minutes daily over 5 days, with re-exposure on day 7. Behavioral tests (SPT, OTT) were conducted on days 1, 5, and 7. Experiment II: fish were exposed to water, a single ketamine immersion (40 mg/L on day 5), or repeated immersions (days 1–5). Tests were conducted on day 7. Experiment III: fish were exposed to water, a single immersion (day 14), or repeated daily immersions (days 1–14) of 40 mg/L ketamine. Testing occurred on day 16. SPT = social preference test; OTT = Open tank test.

### Experiment I

This experiment investigated the behavioral effects of repeated ketamine exposure at different concentrations. A total of 128 adult zebrafish were randomly assigned to one of four treatment groups (n = 32): control (17 females, 15 males), ketamine 10 mg/L (16 females, 16 males), ketamine 20 mg/L (15 females, 17 males), and ketamine 40 mg/L (16 females, 16 males). Missing data in the analysis reflect animal mortality as detailed in the “Statistical analysis” section. Animals received a 20-minute exposure to their assigned treatment once per day from experimental days 1 to 5. For each session, fish were transferred in groups of four to beakers containing 400 mL of the appropriate solution. After the washout period on day 6 (no handling or exposure), a final treatment was administered on day 7. This administration schedule was informed by prior research in mice, which demonstrated behavioral sensitization following five consecutive days of ketamine exposure and a subsequent 48-hours washout period [27]. Following each exposure, animals were returned to their respective home tanks. Behavioral testing included both the social preference test (SPT) and the open tank test (OTT), conducted in sequence on days 1, 5, and 7 immediately after drug exposure. This experiment was designed to test the hypothesis that repeated exposure to ketamine reduces social interaction (as measured by time spent in the interaction zone during the SPT) and increases rotations in the OTT.

### Experiment II

Building on the findings from Experiment I, the 40 mg/L ketamine concentration was selected for further investigation in Experiment II. This experiment evaluated whether behavioral changes induced by ketamine persist after treatment ends, comparing the effects of a single exposure to those of repeated administration. A total of 48 adult zebrafish were randomly assigned to one of three treatment groups (n = 16): control (11 females, 5 males), single exposure (7 females, 9 males), and repeated exposure (7 females, 9 males). Animals in the repeated group received a 20-minute exposure to 40 mg/L ketamine once daily from experimental days 1 to 5. The single exposure group was treated with water from days 1 to 4 and exposed to 40 mg/L ketamine only on day 5. Control animals were exposed to water throughout the same period. For each exposure session, fish were transferred in groups of four to beakers containing 400 mL of the assigned solution. On day 6, animals remained in their home tanks without intervention. Behavioral testing was carried out on day 7, 48 hours after the final exposure. All animals were evaluated in both the SPT and the OTT, in that order. This experiment was designed to test the hypothesis that repeated ketamine exposure induces sustained behavioral alterations as compared to a single exposure.

### Experiment III

Following the results of Experiment II, a new experiment was designed to assess whether a longer exposure period would lead to sustained behavioral effects. A total of 48 adult zebrafish were randomly assigned to one of three treatment groups (n = 16): control (9 females, 7 males), single exposure (5 females, 11 males), and repeated exposure (10 females, 6 males). Missing data in the analysis reflect animal mortality as detailed in the “Statistical analysis” section. Animals in the repeated group received a 20-minute exposure to 40 mg/L ketamine once daily from experimental days 1 to 14. The single exposure group was treated with water from days 1 to 13 and received a single 40 mg/L ketamine exposure on day 14. Control animals were exposed to water throughout the same period. For each exposure session, fish were transferred in groups of four to beakers containing 400 mL of the assigned solution. On day 15, animals remained in their home tanks without intervention. Behavioral testing was carried out on day 16, 48 hours after the final exposure. All animals were evaluated in both the SPT and the OTT, in that order. This experiment was designed to test the hypothesis that repeated ketamine exposure induces sustained behavioral alterations as compared to a single exposure.

### Behavioral tests

The SPT and OTT were conducted sequentially to assess behavioral outcomes. Testing took place during the light phase, between 7:00 a.m. and 1:00 p.m. Animals were recorded from the front view during the SPT and from above during the OTT. Behavioral data were analyzed using ANY-maze® tracking software (version 7.20; Wood Dale, IL, USA). All researchers involved in analyzing the recordings were blinded to group allocation.

### Social preference test

The SPT was used to assess social affiliation in zebrafish [18, 19]. After treatment, each animal was individually placed in a rectangular tank (30 × 10 × 15 cm) positioned between two adjacent tanks (15 × 10 × 13 cm each). One side tank contained a group of 10 conspecifics (social stimulus), while the other was left empty (neutral stimulus). All tanks were filled with water under housing conditions to a height of 10 cm. The location of the social stimulus (left or right) was alternated across subjects to control for side preference. Water in the setup was replaced between tests. For analysis, the test tank was virtually divided lengthwise into three equal zones: the interaction zone (nearest to the social stimulus), the neutral zone (nearest to the empty tank), and a central intermediate zone. Video recordings lasted 7 minutes, consisting of a 2-minute habituation period followed by 5 minutes of data collection. The primary outcome was time spent in the interaction zone. Additional variables included total distance traveled, maximum speed, number of line crossings, percentage of distance traveled in the interaction zone, and time spent immobile.

### Open tank test

The OTT was conducted immediately after the SPT to evaluate locomotor activity and stereotyped swimming behavior [18]. Each fish was placed in the center of a circular white arena (24 cm in diameter, 8 cm high) filled with 2 cm of water under standard conditions. Recordings were captured from an overhead view for a 5-minute session. The arena was refilled with clean water between trials. For behavioral analysis, the arena was virtually divided into two zones: a center zone (12 cm in diameter) and a peripheral zone. The primary variable of interest was the number of rotations, which served as a marker of repetitive or stereotyped movement. Additional outcomes included total distance traveled, maximum speed, time spent in the center zone, percentage of distance traveled in the center, absolute turn angle, and time spent immobile.

### Statistical analysis

For Experiment I, the planned sample size was 48 animals per group (total = 192), based on a power of 0.88 to detect a medium effect size (f = 0.3, α = 0.05). After completing two batches (64 animals each), testing was halted due to consistent results and effect sizes greater than expected. The final sample size was 32 per group (total = 128). A sensitivity analysis indicated that, for post hoc comparisons between two groups (n = 32 each), the design had sufficient power (1-β = 0.88, α = 0.05) to detect a minimum effect size of d = 0.79. Since most of the observed effect sizes exceeded this threshold, the study was adequately powered to detect the effects identified in pairwise comparisons. No outliers were excluded. Six animals died during the experiment: three from the 40 mg/L group (two females during exposure on day 3; one male one hour after exposure on day 4), one male in the control group (bitten by a tankmate on day 5), and one male from each the 10 and 20 mg/L groups (after testing on day 5). For Experiments II and III, the initial plan was 32 animals per group (total = 96), based on a power of 0.94 to detect a large effect (f = 0.4, α = 0.05). After one batch (48 animals), testing was discontinued due to the absence of clear treatment effects and in order to minimize animal use. The final sample size was 16 per group. One male from the control group died during Experiment III; no deaths occurred in Experiment II. Sample sizes were estimated using G*Power (version 3.1; Düsseldorf, Germany). Effect sizes (Cohen’s d) and 95% confidence intervals were calculated using R (version 4.3.1), RStudio (version 2023.12.1+402; Boston, MA, USA), and the meta package (version 8.0-1) [34]. In Experiment I, exploratory subgroup analyses across test days were conducted using a random-effects model. Behavioral data were analyzed in GraphPad Prism (version 10.1.0; Boston, MA, USA). One-way ANOVA followed by Bonferroni post hoc tests was used for parametric outcomes; immobility time was analyzed using the Kruskal–Wallis test followed by Dunn’s correction. Results are reported as mean ± SD for normally distributed outcomes and as median with interquartile range for nonparametric data (as determined by inspecting the Q–Q plots). A p-value below 0.05 was considered statistically significant.

### Deviations from registered protocols

In Experiment I, the preregistered sample size was 48 animals per group (total = 192), but data collection was stopped after 32 animals per group due to consistent results and larger-than-expected effects. The planned analysis involved a two-way ANOVA with treatment group and experimental day as factors. However, because individual animals could not be reliably tracked across days, each day was analyzed independently using one-way ANOVA to avoid inflating the sample size. Additionally, "percent distance in the interaction zone" and "percent distance in the center" were not preregistered outcomes but were included post hoc in Experiment I for consistency with Experiments II and III. In Experiments II and III, although 32 animals per group were planned, testing was halted after one batch (16 per group) due to a lack of apparent treatment effects and to reduce unnecessary animal use, in accordance with the 3Rs principle [35]. Across all experiments, immobility time was analyzed using Kruskal–Wallis followed by Dunn’s test instead of ANOVA, as this outcome violated parametric assumptions.

## RESULTS

### Experiment I

Ketamine administration led to consistent, concentration-dependent behavioral changes across tests. In the SPT, all concentrations reduced time spent in the interaction zone and percent distance in the interaction zone, indicating decreased social preference. At the same time, ketamine increased total distance traveled, maximum speed, and number of line crossings, reflecting a pattern of hyperlocomotion. In the OTT, ketamine also increased number of rotations, distance traveled, and maximum speed, especially at higher concentrations, while the lower concentration increased time spent in the center and percent distance in the center, an anxiolytic-like response typically interpreted as reduced thigmotaxis in an unfamiliar environment. Subgroup analyses revealed no significant differences across testing days for any outcome, suggesting that repeated ketamine exposure did not result in behavioral sensitization. The following sections provide a more detailed account of these results.

### Social preference test

Figure 2 shows the effects of different ketamine concentrations (10, 20, and 40 mg/L) in the SPT across three test sessions (days 1, 5, and 7). A significant effect of treatment was observed on each test day for the primary outcome, with ketamine reducing time spent in the interaction zone (day 1: F(3,124) = 32.96, p < 0.0001; day 5: F(3,120) = 13.80, p < 0.0001; day 7: F(3,118) = 12.73, p < 0.0001; Figure 2A). Bonferroni post hoc comparisons showed that all ketamine concentrations significantly reduced social preference compared to the control group on all three days. Effect sizes are presented in Figure 2B. On day 1, effect sizes were directly proportional to ketamine concentration, indicating a concentration-dependent increase in effect magnitude: −1.21 (95% CI [-1.75, −0.68]) for 10 mg/L, −1.76 [-2.33, −1.18] for 20 mg/L, and −2.87 [-3.57, −2.16] for 40 mg/L. This pattern persisted on day 5 (−0.57 [-1.07, −0.06] for 10 mg/L; −0.90 [-1.42, - 0.38] for 20 mg/L; −1.71 [-2.31, −1.12] for 40 mg/L) and day 7 (−0.76 [-1.27, −0.24] for 10 mg/L; −0.84 [-1.36, −0.32] for 20 mg/L; −1.71 [-2.31, −1.12] for 40 mg/L). No significant differences were found across testing days in the subgroup analysis (p = 0.2760).

**Figure 2.**
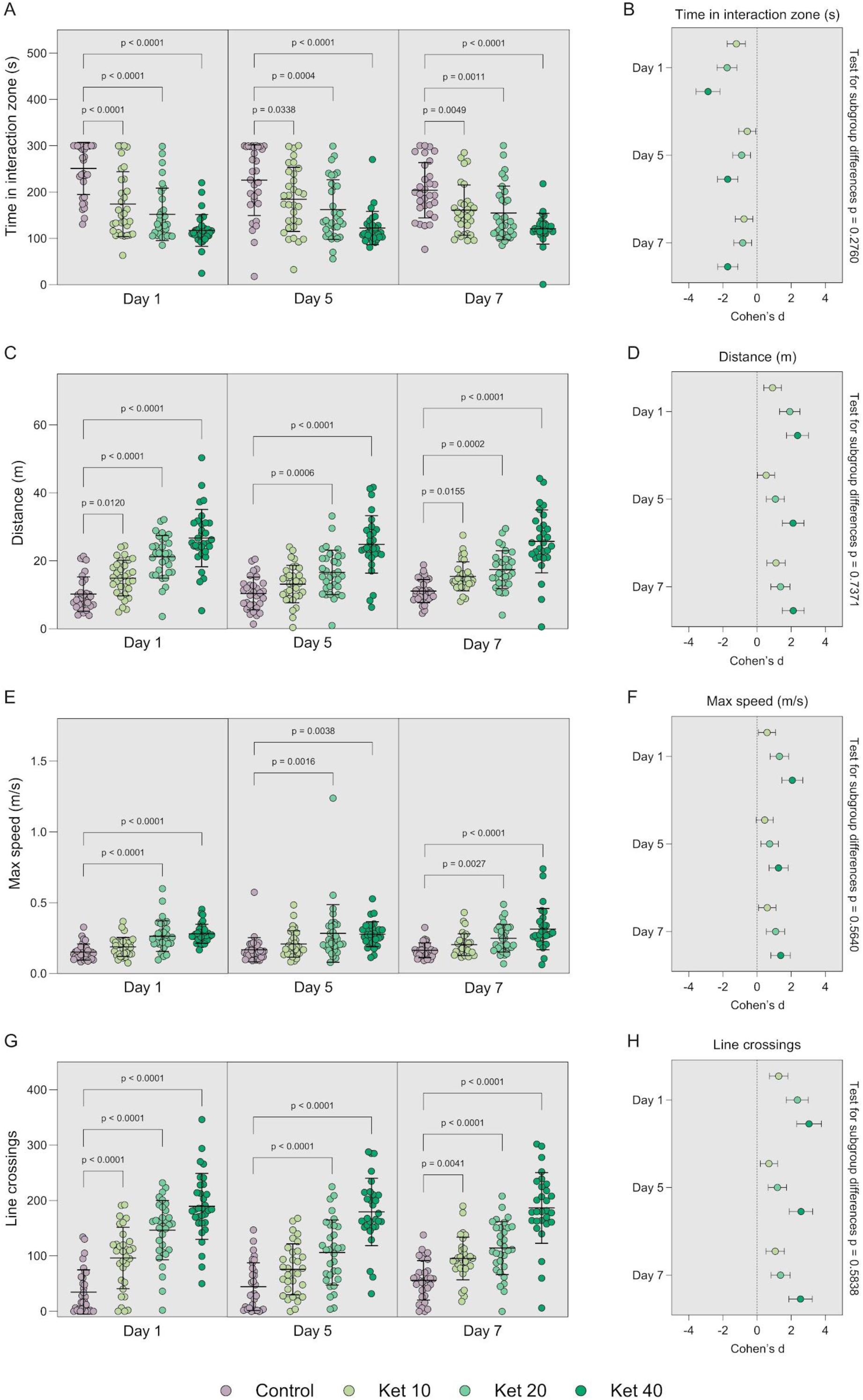
Effects of exposure to different concentrations of ketamine on the behavior of adult zebrafish in the social preference test. (A) Time spent in the interaction zone, (C) total distance traveled, (E) maximum speed, and (G) number of line crossings, measured on days 1, 5, and 7. (B, D, F, H) Corresponding effect sizes (Cohen’s d) for each outcome, with test for subgroup differences across assessment days. One-way ANOVA followed by Bonferroni post hoc test. Data are presented as mean ± SD. Effect sizes are expressed as Cohen’s d with 95% confidence intervals. Subgroup differences tested using a random-effects model. Sample sizes: day 1 – Control = 32, Ket 10 = 32, Ket 20 = 32, Ket 40 = 32; day 5 – Control = 31, Ket 10 = 32, Ket 20 = 32, Ket 40 = 29; day 7 – Control = 31, Ket 10 = 31, Ket 20 = 31, Ket 40 = 29. Ket 10 = ketamine 10 mg/L; Ket 20 = ketamine 20 mg/L; Ket 40 = ketamine 40 mg/L.

Ketamine also significantly increased total distance traveled during the test sessions (Figure 2C). An effect of treatment was detected on all days (day 1: F(3,124) = 40.58, p < 0.0001; day 5: F(3,120) = 28.25, p < 0.0001; day 7: F(3,118) = 31.25, p < 0.0001). Post hoc tests indicated significant increases at all concentrations on day 1, at 20 and 40 mg/L on day 5, and again for all groups on day 7. Effect sizes are shown in Figure 2D. On each day, effect sizes increased with concentration: 0.91 [0.40, 1.43] for 10 mg/L, 1.92 [1.32, 2.51] for 20 mg/L, and 2.37 [1.73, 3.02] for 40 mg/L on day 1; 0.53 [0.03, 1.04] for 10 mg/L, 1.08 [0.55, 1.61] for 20 mg/L, and 2.12 [1.48, 2.75] for 40 mg/L on day 5; and 1.11 [0.58, 1.65] for 10 mg/L, 1.37 [0.82, 1.93] for 20 mg/L, and 2.12 [1.49, 2.76] for 40 mg/L on day 7. No significant differences were found across testing days in the subgroup analysis (p = 0.7371).

A significant main effect of treatment was also observed for maximum swimming speed (Figure 2E), with increases detected on each test day (day 1: F(3,124) = 20.31, p < 0.0001; day 5: F(3,120) = 5.82, p = 0.001; day 7: F(3,118) = 12.81, p < 0.0001). Post hoc analysis showed that only the 20 and 40 mg/L groups significantly differed from controls across all days, with no significant effects at 10 mg/L. As shown in Figure 2F, effect sizes increased with concentration: 0.59 [0.09, 1.09] for 10 mg/L, 1.31 [0.77, 1.85] for 20 mg/L, and 2.07 [1.46, 2.68] for 40 mg/L on day 1; 0.46 [-0.04, 0.96] for 10 mg/L, 0.74 [0.22, 1.25] for 20 mg/L, and 1.26 [0.71, 1.82] for 40 mg/L on day 5; and 0.60 [0.09, 1.11] for 10 mg/L, 1.09 [0.55, 1.62] for 20 mg/L, and 1.38 [0.82, 1.95] for 40 mg/L on day 7. No significant differences were found across testing days in the subgroup analysis (p = 0.5640).

Line crossings, another index of locomotor activity, was also increased following ketamine exposure (Figure 2G) on all days (day 1: F(3,124) = 51.21, p < 0.0001; day 5: F(3,120) = 35.98, p < 0.0001; day 7: F(3,118) = 39.69, p < 0.0001). All ketamine groups differed from controls on days 1 and 7, while only the 20 and 40 mg/L concentrations were significant on day 5. Effect sizes are shown in Figure 2H. On each test day, values increased with concentration: 1.27 [0.73, 1.81] for 10 mg/L, 2.37 [1.72, 3.01] for 20 mg/L, and 3.05 [2.32, 3.77] for 40 mg/L on day 1; 0.70 [0.19, 1.21] for 10 mg/L, 1.20 [0.66, 1.73] for 20 mg/L, and 2.57 [1.88, 3.27] for 40 mg/L on day 5; and 1.06 [0.53, 1.60] for 10 mg/L, 1.38 [0.83, 1.94] for 20 mg/L, and 2.55 [1.86, 3.24] for 40 mg/L on day 7. No significant differences were found across testing days in the subgroup analysis (p = 0.5838).

Treatment also significantly decreased the percent distance traveled within the interaction zone (Supplementary Figure 1A). An effect was observed on all test days (day 1: F(3,124) = 28.64, p < 0.0001; day 5: F(3,120) = 15.12, p < 0.0001; day 7: F(3,118) = 7.67, p = 0.0001). Post hoc analysis confirmed that all concentrations significantly reduced this metric compared to controls on each day. Effect sizes are shown in Supplementary Figure 1B. Effect magnitudes increased with concentration, reflected by more negative values: −1.13 [-1.66, −0.60] for 10 mg/L, −1.67 [-2.24, −1.10] for 20 mg/L, and −2.44 [-3.10, −1.79] for 40 mg/L on day 1; −0.75 [-1.26, −0.24] for 10 mg/L, −1.10 [-1.64, −0.57] for 20 mg/L, and −1.75 [-2.35, - 1.15] for 40 mg/L on day 5; and −0.56 [-1.07, −0.06] for 10 mg/L, −0.56 [-1.07, −0.06] for 20 mg/L, and −1.22 [-1.77, −0.66] for 40 mg/L on day 7. No significant differences were found across testing days in the subgroup analysis (p = 0.0770).

No significant effect of ketamine was observed on time spent immobile during the social preference test (Supplementary Figure 1C). Kruskal–Wallis tests yielded non-significant results on all test days (day 1: H = 7.09, p = 0.0692; day 5: H = 3.02, p = 0.3879; day 7: H = 1.77, p = 0.6223). Effect sizes are presented in Supplementary Figure 1D. On day 1, values were −0.36 [-0.85, 0.14] for 10 mg/L, −0.30 [-0.79, 0.19] for 20 mg/L, and −0.17 [-0.66, 0.32] for 40 mg/L. Day 5 values were 0.00 [-0.49, 0.50] for 10 mg/L, −0.30 [-0.80, 0.19] for 20 mg/L, and −0.43 [-0.95, 0.08] for 40 mg/L. On day 7, effect sizes were −0.39 [-0.90, 0.11] for 10 mg/L, −0.42 [-0.92, 0.08] for 20 mg/L, and 0.15 [-0.36, 0.65] for 40 mg/L. No significant differences were found across testing days in the subgroup analysis (p = 0.9385).

### Open tank test

Figure 3 summarizes the results of the OTT, assessing multiple behavioral outcomes in response to different ketamine concentrations (10, 20, and 40 mg/L) across three sessions (days 1, 5, and 7). For the primary outcome, number of rotations, a significant effect of treatment was observed on day 1 (F(3,124) = 5.32, p = 0.0018) and day 5 (F(3,120) = 9.61, p < 0.0001), but not on day 7 (F(3,118) = 1.73, p = 0.1637; Figure 3A). Post hoc comparisons indicated significant increases in rotations for animals treated with 20 and 40 mg/L on day 1, and for 40 mg/L on day 5. Effect sizes and 95% confidence intervals are shown in Figure 3B. On day 1, values were: 0.15 [-0.34, 0.64] for 10 mg/L, 0.68 [0.18, 1.19] for 20 mg/L, and 0.94 [0.42, 1.46] for 40 mg/L. Day 5 showed a similar trend, with effect sizes of 0.22 [-0.27, 0.72] for 10 mg/L, 0.38 [-0.12, 0.87] for 20 mg/L, and 1.15 [0.60, 1.70] for 40 mg/L. On day 7, effect sizes were 0.21 [-0.29, 0.71] for 10 mg/L, −0.02 [-0.52, 0.47] for 20 mg/L, and 0.63 [0.11, 1.15] for 40 mg/L. Effect sizes increased with ketamine concentration across all days. No significant differences were found across testing days in the subgroup analysis (p = 0.4911).

**Figure 3.**
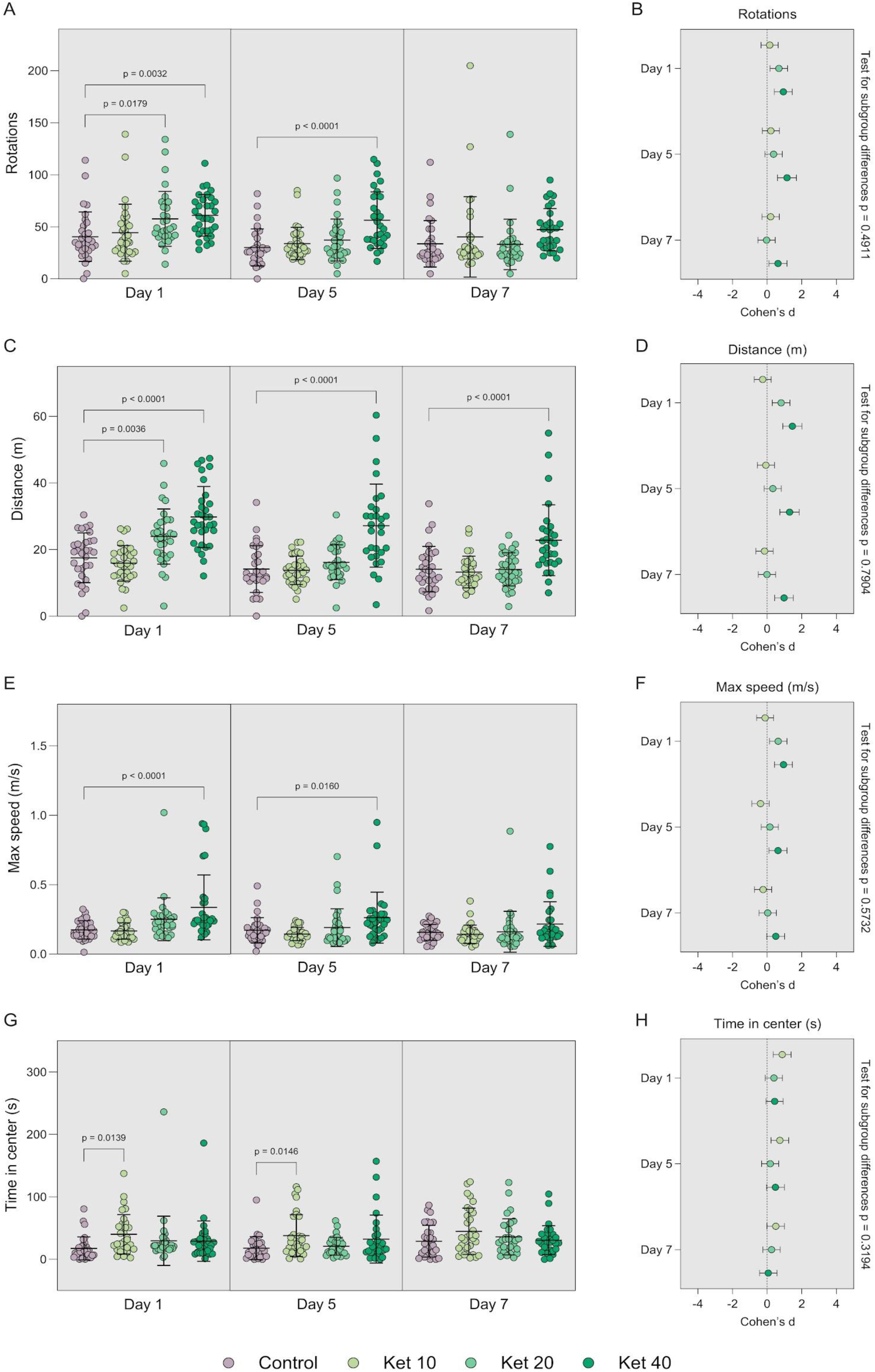
Effects of exposure to different concentrations of ketamine on the behavior of adult zebrafish in the open tank test. (A) Number of rotations, (C) total distance traveled, (E) maximum speed, and (G) time spent in the center zone, measured on days 1, 5, and 7. (B, D, F, H) Corresponding effect sizes (Cohen’s d) for each outcome, with test for subgroup differences across assessment days. One-way ANOVA followed by Bonferroni post hoc test. Data are presented as mean ± SD. Effect sizes are expressed as Cohen’s d with 95% confidence intervals. Subgroup differences tested using a random-effects model. Sample sizes: day 1 – Control = 32, Ket 10 = 32, Ket 20 = 32, Ket 40 = 32; day 5 – Control = 31, Ket 10 = 32, Ket 20 = 32, Ket 40 = 29; day 7 – Control = 31, Ket 10 = 31, Ket 20 = 31, Ket 40 = 29. Ket 10 = ketamine 10 mg/L; Ket 20 = ketamine 20 mg/L; Ket 40 = ketamine 40 mg/L.

For distance traveled, a significant effect of treatment was observed on day 1 (F(3,124) = 21.81, p < 0.0001), day 5 (F(3,120) = 19.72, p < 0.0001), and day 7 (F(3,118) = 11.86, p < 0.0001; Figure 3C). Post hoc comparisons revealed significant increases in distance for animals treated with 20 and 40 mg/L on day 1, and only 40 mg/L on days 5 and 7. Effect sizes and 95% confidence intervals are displayed in Figure 3D. On day 1, values were: −0.25 [-0.74, 0.24] for 10 mg/L, 0.81 [0.30, 1.32] for 20 mg/L, and 1.46 [0.91, 2.02] for 40 mg/L. On day 5, effect sizes were −0.07 [-0.56, 0.43] for 10 mg/L, 0.33 [-0.17, 0.83] for 20 mg/L, and 1.30 [0.74, 1.86] for 40 mg/L. On day 7, values were −0.14 [-0.64, 0.36] for 10 mg/L, −0.01 [-0.50, 0.49] for 20 mg/L, and 0.98 [0.44, 1.52] for 40 mg/L. The magnitude of the effect sizes increased with ketamine concentration on each day. No significant differences were found across testing days in the subgroup analysis (p = 0.7904).

For maximum speed, a significant effect of treatment was found on day 1 (F(3,124) = 9.34, p < 0.0001) and day 5 (F(3,120) = 4.97, p = 0.0028), but not on day 7 (F(3,118) = 2.31, p = 0.0799; Figure 3E). Post hoc analyses revealed significant increases in maximum speed only for the 40 mg/L group on days 1 and 5. Effect sizes and 95% confidence intervals are presented in Figure 3F. On day 1, values were: −0.11 [−0.60, 0.38] for 10 mg/L, 0.65 [0.14, 1.15] for 20 mg/L, and 0.95 [0.43, 1.46] for 40 mg/L. On day 5, values were: −0.38 [-0.87, 0.12] for 10 mg/L, 0.16 [-0.33, 0.66] for 20 mg/L, and 0.63 [0.11, 1.15] for 40 mg/L. On day 7, effect sizes were −0.23 [-0.73, 0.27] for 10 mg/L, 0.04 [-0.46, 0.54] for 20 mg/L, and 0.50 [-0.01, 1.02] for 40 mg/L. No significant differences were found across testing days in the subgroup analysis (p = 0.5732).

For time spent in the center, a significant effect of treatment was found on day 1 (F(3,124) = 2.78, p = 0.0439) and day 5 (F(3,120) = 3.65, p = 0.0146), but not on day 7 (F(3,118) = 1.84, p = 0.1441; Figure 3G). Post hoc comparisons indicated significantly increased center time only for animals treated with 10 mg/L on days 1 and 5. Effect sizes and 95% confidence intervals are provided in Figure 3H. On day 1, values were: 0.87 [0.36, 1.39] for 10 mg/L, 0.40 [-0.10, 0.89] for 20 mg/L, and 0.44 [-0.06, 0.93] for 40 mg/L. On day 5, effect sizes were 0.74 [0.23, 1.25] for 10 mg/L, 0.18 [-0.31, 0.68] for 20 mg/L, and 0.49 [-0.03, 1.00] for 40 mg/L. By day 7, effects diminished: 0.50 [-0.01, 1.01] for 10 mg/L, 0.26 [-0.24, 0.76] for 20 mg/L, and 0.07 [-0.43, 0.58] for 40 mg/L. The magnitude of the effect was highest for 10 mg/L across all days. Increased center time is interpreted as an anxiolytic effect, and was only observed at the lowest concentration. No significant differences were found across testing days in the subgroup analysis (p = 0.3194).

For percent distance in the center, a significant effect of treatment was detected on day 1 (F(3,124) = 6.01, p = 0.0007) and day 5 (F(3,120) = 3.18, p = 0.0266), but not on day 7 (F(3,118) = 1.57, p = 0.2015; Supplementary Figure 2A). Post hoc analysis revealed significant increases only for animals treated with 10 mg/L on days 1 and 5. Effect sizes and 95% confidence intervals are shown in Supplementary Figure 2B. On day 1, effect sizes were: 1.03 [0.51, 1.55] for 10 mg/L, 0.59 [0.09, 1.09] for 20 mg/L, and 0.45 [-0.04, 0.95] for 40 mg/L. Day 5 values were: 0.71 [0.20, 1.22] for 10 mg/L, 0.18 [-0.32, 0.67] for 20 mg/L, and 0.43 [-0.08, 0.94] for 40 mg/L. By day 7, effect sizes decreased: 0.44 [-0.06, 0.95] for 10 mg/L, 0.25 [-0.25, 0.75] for 20 mg/L, and 0.04 [-0.47, 0.55] for 40 mg/L. Increased percent distance in center is interpreted as anxiolytic-like behavior, observed only at the lowest ketamine concentration. No significant differences were found across testing days in the subgroup analysis (p = 0.1010).

For absolute turn angle, no significant effects of treatment were observed on day 1 (F(3,124) = 0.62, p = 0.6036), day 5 (F(3,120) = 1.11, p = 0.3485), or day 7 (F(3,118) = 0.56, p = 0.6395; Supplementary Figure 2C). No group significantly differed from controls on any day. Effect sizes are shown in Supplementary Figure 2D. On day 1: −0.25 [-0.74, 0.24] for 10 mg/L, 0.03 [-0.46, 0.52] for 20 mg/L, and −0.07 [-0.56, 0.42] for 40 mg/L. On day 5: −0.02 [-0.51, 0.48] for 10 mg/L, −0.37 [-0.86, 0.13] for 20 mg/L, and −0.15 [-0.66, 0.36] for 40 mg/L. On day 7: −0.08 [-0.58, 0.41] for 10 mg/L, −0.27 [-0.77, 0.23] for 20 mg/L, and −0.02 [-0.53, 0.48] for 40 mg/L. No significant differences were found across testing days in the subgroup analysis (p = 0.8217).

No significant effect of ketamine was observed on time spent immobile during the open tank test on any of the three experimental days (day 1: H = 7.68, p = 0.053; day 5: H = 6.73, p = 0.0812; day 7: H = 3.63, p = 0.3038; Supplementary Figure 2E). Effect sizes and 95% confidence intervals are shown in Supplementary Figure 2F. On day 1: −0.24 [-0.73, 0.25] for 10 mg/L, −0.29 [-0.78, 0.21] for 20 mg/L, and - 0.36 [-0.86, 0.13] for 40 mg/L. On day 5: −0.27 [-0.77, 0.22] for 10 mg/L, −0.29 [-0.79, 0.21] for 20 mg/L, and −0.48 [-1.00, 0.03] for 40 mg/L. On day 7: −0.43 [-0.93, 0.08] for 10 mg/L, −0.30 [-0.80, 0.20] for 20 mg/L, and −0.44 [-0.95, 0.07] for 40 mg/L. No significant differences were found across testing days in the subgroup analysis (p = 0.2612).

### Experiment II

Ketamine administration at 40 mg/L failed to produce sustained behavioral alterations following either acute or repeated exposure. In the SPT, neither a single administration on day 5 nor repeated treatment over 5 consecutive days affected time spent in the interaction zone, percent distance in the interaction zone, or locomotor-related measures such as distance traveled, line crossings, maximum speed, or time immobile. Likewise, in the OTT, no significant effects were observed for number of rotations, distance traveled, time spent in the center, percent distance in the center, absolute turn angle, or time immobile. A significant reduction in maximum speed was detected only in the group receiving repeated administration, suggesting a limited and transient effect. Overall, the absence of robust behavioral changes after a 48-hour washout indicates that neither acute nor repeated ketamine exposure at this concentration produced sustained effects. The following sections provide a more detailed account of these results.

### Social preference test

Figure 4 shows the effects of single and repeated ketamine exposure (40 mg/L) on behavior in the SPT. In this protocol, ketamine was administered daily for five consecutive days, and behavioral testing was conducted 48 hours after the final concentration. For the primary outcome, time spent in the interaction zone, no significant effect of treatment was observed (F(2,45) = 0.40, p = 0.6701; Figure 4A). Effect sizes are shown in Figure 4G: −0.32 [-1.02, 0.37] for the single exposure group and −0.11 [-0.81, 0.58] for the repeated group. For distance traveled, there was also no significant treatment effect (F(2,45) = 0.60, p = 0.5524; Figure 4B), with effect sizes of 0.25 [-0.45, 0.94] and 0.42 [-0.28, 1.12] for single and repeated exposures, respectively. No effect of treatment was detected for maximum speed (F(2,45) = 0.22, p = 0.8066; Figure 4C), and effect sizes were 0.13 [-0.56, 0.83] for single and −0.10 [-0.79, 0.59] for repeated exposure. Similarly, no significant effect was found for the number of line crossings (F(2,45) = 0.09, p = 0.9137; Figure 4D), with effect sizes of 0.11 [-0.59, 0.80] and −0.05 [-0.74, 0.65], respectively. For percent distance in the interaction zone (F(2,45) = 0.09, p = 0.9172; Figure 4E), results were non-significant, with effect sizes of −0.10 [-0.80, 0.59] for single and 0.05 [-0.65, 0.74] for repeated exposure. Finally, for time immobile, no significant difference was found (H = 0.325, p = 0.8501; Figure 4F), and the effect sizes were 0.31 [-0.39, 1.01] for single and 0.12 [-0.58, 0.81] for repeated exposure.

**Figure 4.**
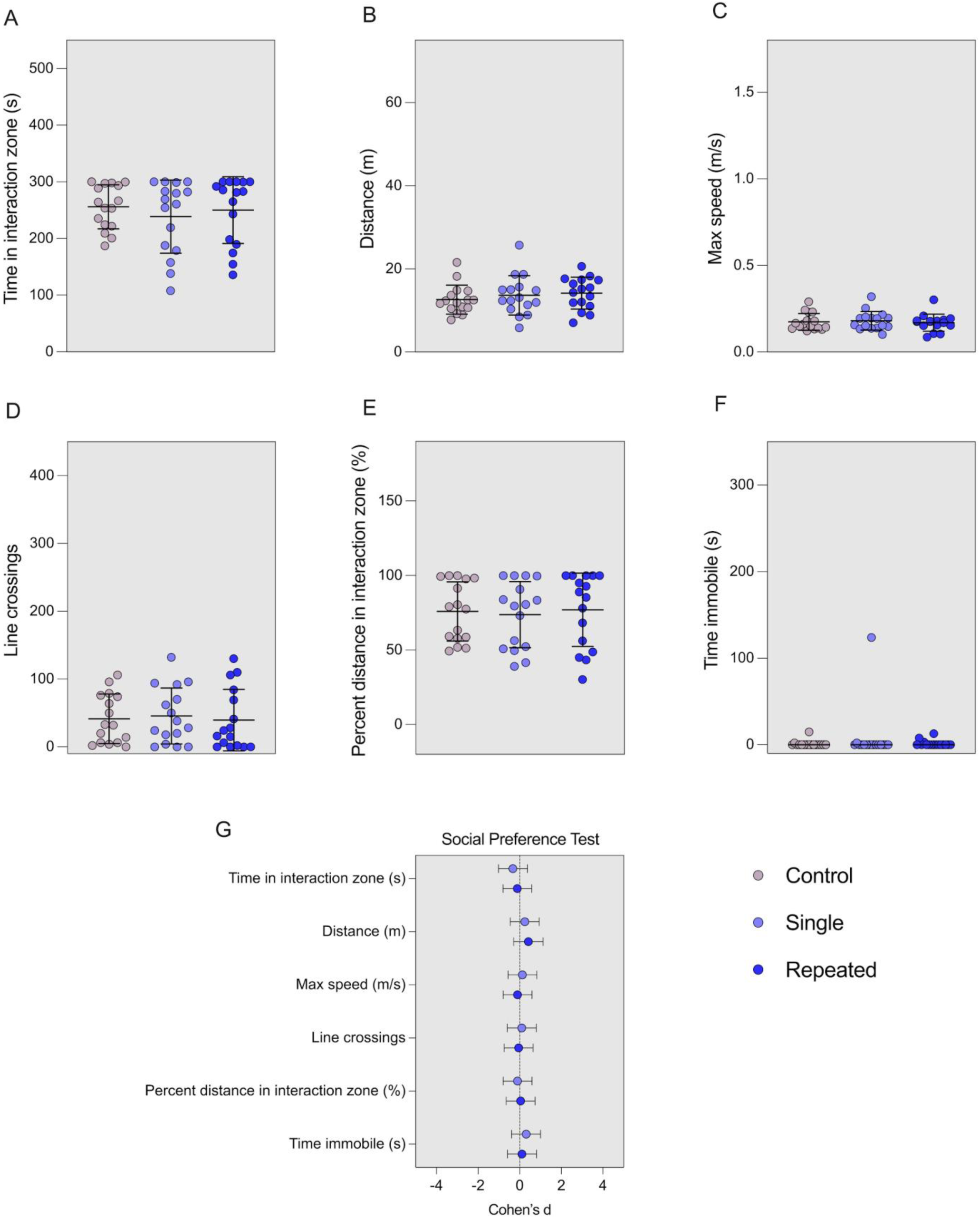
Effects of single (day 5) or repeated (days 1–5) ketamine exposures on the behavior of adult zebrafish in the social preference test. (A) Time spent in the interaction zone, (B) total distance traveled, (C) maximum speed, (D) number of line crossings, (E) percent of distance traveled in the interaction zone, and (F) time spent immobile. (G) Effect sizes (Cohen’s d) for the single and repeated exposure groups across the outcomes assessed. One-way ANOVA followed by Bonferroni post hoc test was used for A–E. Kruskal–Wallis test followed by Dunn’s post hoc was used for F. Data are presented as mean ± SD for A– E, and as median with interquartile range for F. Effect sizes are expressed as Cohen’s d with 95% confidence intervals. Sample sizes: Control = 16, Single = 16, Repeated = 16. Single = single ketamine exposure (40 mg/L) on day 5; Repeated = daily ketamine exposures (40 mg/L) from day 1 to 5.

### Open tank test

Figure 5 presents the results of the OTT conducted after single or repeated ketamine exposure (40 mg/L). In this protocol, ketamine was administered daily for five consecutive days, and behavioral testing was conducted 48 hours after the final concentration. For the primary outcome, number of rotations, no significant differences were detected between groups (F(2,45) = 0.52 p = 0.5994; Figure 5A), with effect sizes of 0.12 [-0.57, 0.82] and 0.33 [-0.37, 1.03] for the single and repeated groups, respectively. For distance traveled, results were not significant (F(2,45) = 0.36, p = 0.6991; Figure 5B), and the effect sizes were −0.11 [-0.80, 0.58] for single and 0.18 [-0.51, 0.88] for repeated exposure. Maximum speed showed a significant treatment effect (F(2,45) = 3.61, p = 0.0352; Figure 5C), and post hoc comparisons indicated a significant decrease in the repeated ketamine group relative to control. Effect sizes were −0.71 [-1.42, 0.01] for single and −0.82 [-1.55, −0.10] for repeated exposure. For time spent in the center, no significant effect was found (F(2,45) = 1.51, p = 0.2316; Figure 5D), and the corresponding effect sizes were 0.45 [−0.25, 1.16] and 0.13 [-0.56, 0.83]. Likewise, no treatment effect was observed for percent distance in the center (F(2,45) = 1.39, p = 0.2608; Figure 5E), with effect sizes of 0.39 [-0.31, 1.09] and −0.23 [-0.92, 0.47], respectively. No differences were found for absolute turn angle (F(2,45) = 0.35, p = 0.7091; Figure 5F), with effect sizes of 0.03 [-0.66, 0.73] and 0.29 [-0.41, 0.99]. Lastly, for time immobile, no significant difference was observed (H = 0.302, p = 0.8597; Figure 5G), and effect sizes were 0.00 [-0.69, 0.69] and −0.36 [-1.06, 0.34] for single and repeated exposure, respectively.

**Figure 5.**
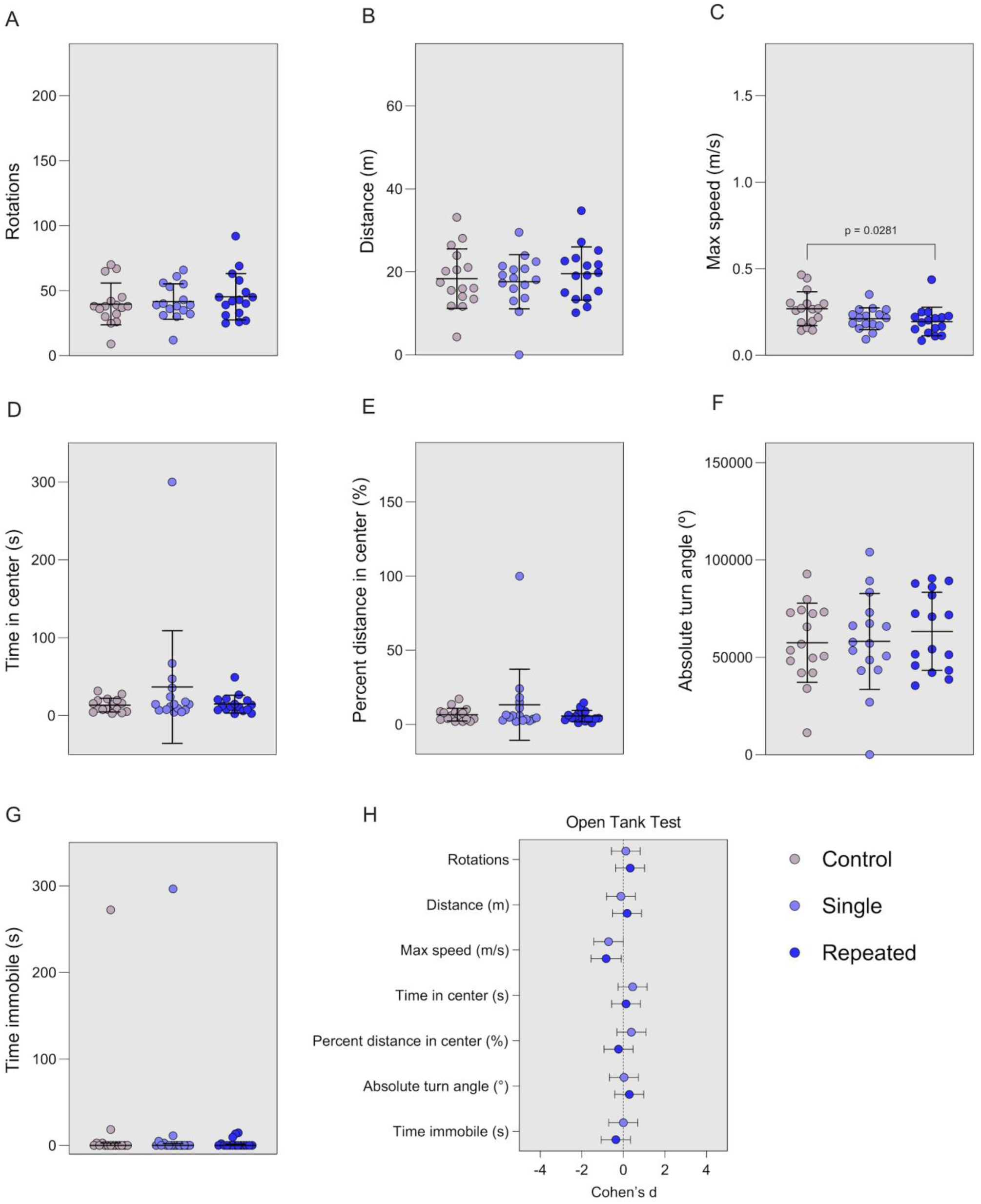
Effects of single (day 5) or repeated (days 1–5) ketamine exposures on the behavior of adult zebrafish in the open tank test. (A) Number of rotations, (B) total distance traveled, (C) maximum speed, (D) time spent in the center zone, (E) percent of distance traveled in the center zone, (F) absolute turn angle, and (G) time spent immobile. (H) Effect sizes (Cohen’s d) for the single and repeated exposure groups across the outcomes assessed. One-way ANOVA followed by Bonferroni post hoc test was used for A–F and G. Kruskal–Wallis test followed by Dunn’s post hoc was used for G. Data are presented as mean ± SD for A–F and G, and as median with interquartile range for G. Effect sizes are expressed as Cohen’s d with 95% confidence intervals. Sample sizes: Control = 16, Single = 16, Repeated = 16. Single = single ketamine exposure (40 mg/L) on day 5; Repeated = daily ketamine exposures (40 mg/L) from day 1 to 5.

### Experiment III

Ketamine administration at 40 mg/L did not produce sustained behavioral effects following either single exposure on day 14 or repeated administration over 14 consecutive days. In the SPT, no differences were observed in time spent in the interaction zone, distance traveled, maximum speed, line crossings, or percent distance in the interaction zone 48 hours after treatment, with the exception of increased time immobile in the repeated group. In the OTT, no treatment effects were detected across any of the outcomes, including number of rotations, distance traveled, maximum speed, time in the center, percent distance in the center, absolute turn angle, and time immobile. Together, these findings indicate a lack of persistent behavioral alterations after ketamine exposure under this protocol. The following sections provide a more detailed account of these results.

### Social preference test

Figure 6 shows the effects of single and repeated ketamine exposure (40 mg/L) on behavior in the SPT. In this protocol, ketamine was administered daily for 14 consecutive days, and behavioral testing was conducted 48 hours after the final concentration. For the primary outcome, time spent in the interaction zone, no significant effect of treatment was observed (F(2,44) = 1.56, p = 0.2214; Figure 6A). Effect sizes were 0.48 [-0.23, 1.20] for the single exposure group and 0.54 [-0.17, 1.26] for the repeated group (Figure 6G). For distance traveled, no significant differences were found (F(2,44) = 1.95, p = 0.1539; Figure 6B), with effect sizes of −0.57 [-1.29, 0.15] and −0.59 [-1.31, 0.14] for the single and repeated groups, respectively. Similarly, no significant treatment effect was observed for maximum speed (F(2,44) = 0.10, p = 0.9049; Figure 6C), with effect sizes of −0.12 [-0.83, 0.58] for single and 0.02 [-0.68, 0.73] for repeated exposure. Line crossings did not differ significantly across groups (F(2,44) = 1.86, p = 0.1678; Figure 6D), with effect sizes of −0.53 [-1.25, 0.19] for single and −0.58 [-1.30, 0.14] for repeated treatment. For percent distance in the interaction zone, results were non-significant (F(2,44) = 1.09, p = 0.3444; Figure 6E), and effect sizes were 0.33 [-0.38, 1.04] and 0.50 [-0.22, 1.22], respectively. Finally, for time immobile, a significant treatment effect was observed (H = 6.788, p = 0.0336; Figure 6F), with post hoc comparisons indicating significantly greater immobility in the repeated group compared to controls. The effect sizes were −0.01 [-0.71, 0.69] for single and 0.39 [-0.32, 1.11] for repeated ketamine exposure (Figure 6G), suggesting an increase in immobility following chronic treatment.

**Figure 6.**
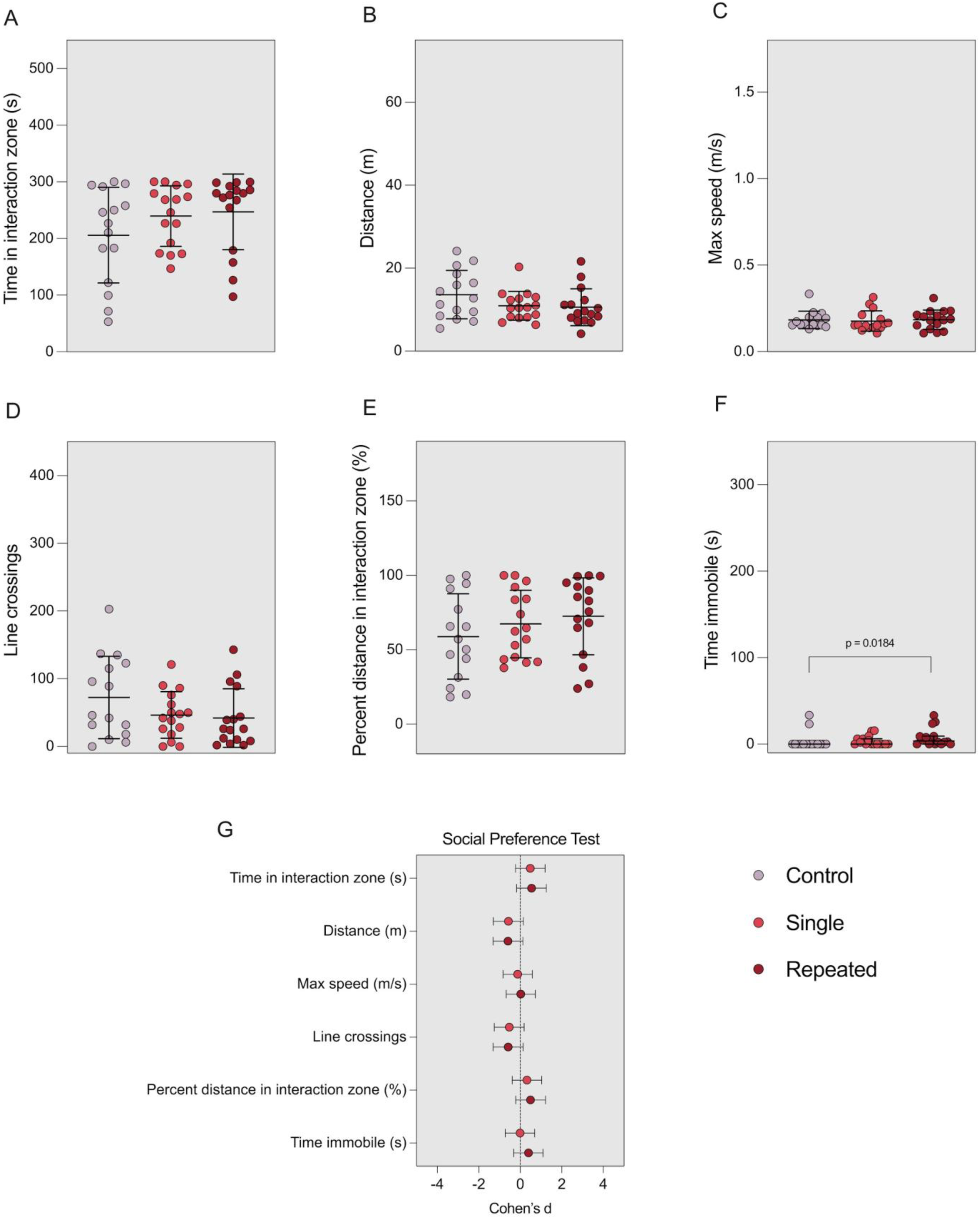
Effects of single (day 14) or repeated (days 1–14) ketamine exposures on the behavior of adult zebrafish in the social preference test. (A) Time spent in the interaction zone, (B) total distance traveled, (C) maximum speed, (D) number of line crossings, (E) percent of distance traveled in the interaction zone, and (F) time spent immobile. (G) Effect sizes (Cohen’s d) for the single and repeated exposure groups across the outcomes assessed. One-way ANOVA followed by Bonferroni post hoc test was used for A–E. Kruskal–Wallis test followed by Dunn’s post hoc was used for F. Data are presented as mean ± SD for A– E, and as median with interquartile range for F. Effect sizes are expressed as Cohen’s d with 95% confidence intervals. Sample sizes: Control = 15, Single = 16, Repeated = 16. Single = single ketamine exposure (40 mg/L) on day 14; Repeated = daily ketamine exposures (40 mg/L) from day 1 to 14.

### Open tank test

Figure 7 presents the results of the OTT following single or repeated ketamine exposure (40 mg/L) over 14 days, with testing conducted after a 48-hour washout. For the primary outcome, number of rotations, no significant treatment effects were found (F(2,44) = 0.06, p = 0.9458; Figure 7A), and effect sizes were 0.11 [-0.60, 0.81] and 0.05 [-0.66, 0.75] for single and repeated groups, respectively (Figure 7H). No significant differences were observed for distance traveled (F(2,44) = 0.01, p = 0.9939; Figure 7B), with effect sizes of 0.04 [-0.67, 0.74] and 0.02 [-0.68, 0.73]. Maximum speed also did not differ across groups (F(2,44) = 2.00, p = 0.1475; Figure 7C), and effect sizes were −0.71 [-1.43, 0.02] for single and −0.49 [-1.21, 0.22] for repeated exposure. For time spent in the center of the tank, no treatment effects were observed (F(2,44) = 0.09, p = 0.9146; Figure 7D), and effect sizes were 0.14 [-0.57, 0.84] and 0.12 [-0.58, 0.83]. Percent distance in the center also showed no significant differences (F(2,44) = 0.62, p = 0.5407; Figure 7E), with effect sizes of −0.25 [-0.96, 0.46] and −0.41 [-1.12, 0.30], respectively. No significant treatment effect was found for absolute turn angle (F(2,44) = 0.46, p = 0.6368; Figure 7F), with effect sizes of 0.27 [−0.44, 0.98] and 0.33 [-0.38, 1.04]. Finally, time immobile did not differ significantly across groups (H = 2.585, p = 0.2746; Figure 7G), and effect sizes were −0.45 [-1.16, 0.27] for single and −0.37 [-1.09, 0.34] for repeated exposure (Figure 7H).

**Figure 7.**
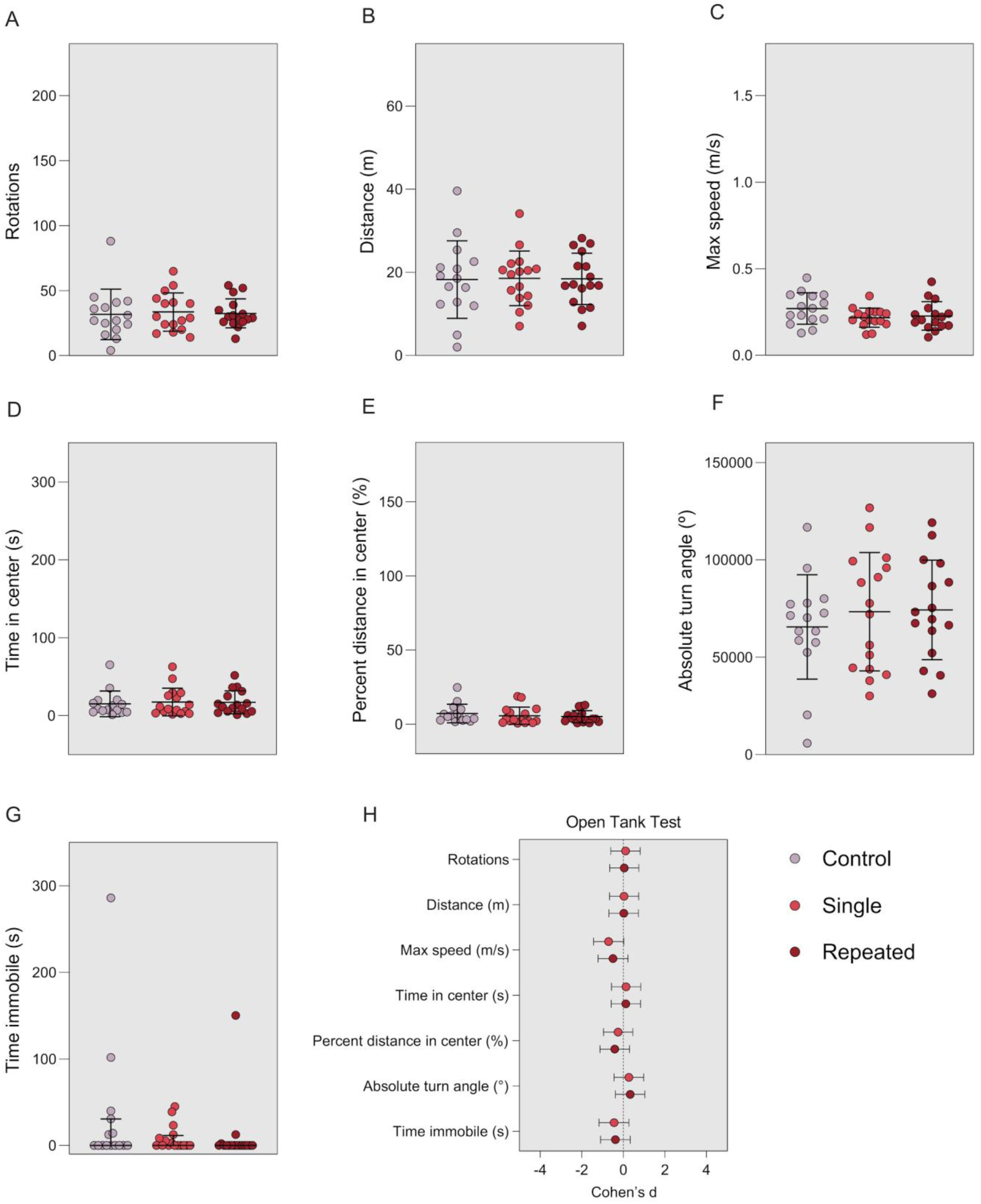
Effects of single (day 14) or repeated (days 1–14) ketamine exposures on the behavior of adult zebrafish in the open tank test. (A) Number of rotations, (B) total distance traveled, (C) maximum speed, (D) time spent in the center zone, (E) percent of distance traveled in the center zone, (F) absolute turn angle, and (G) time spent immobile. (H) Effect sizes (Cohen’s d) for the single and repeated exposure groups across the outcomes assessed. One-way ANOVA followed by Bonferroni post hoc test was used for A–F. Kruskal–Wallis test followed by Dunn’s post hoc was used for G. Data are presented as mean ± SD for A–F, and as median with interquartile range for G. Effect sizes are expressed as Cohen’s d with 95% confidence intervals. Sample sizes: Control = 15, Single = 16, Repeated = 16. Single = single ketamine exposure (40 mg/L) on day 14; Repeated = daily ketamine exposures (40 mg/L) from day 1 to 14.

## DISCUSSION

The present findings reinforce the potential of zebrafish as a translational model for investigating schizophrenia pathophysiology and screening early-stage antipsychotic candidates, while also showing relevant differences following repeated ketamine exposure as compared to rodents. Literature reviews highlight discrepancies regarding the effects of NMDA antagonists on social and locomotor parameters in zebrafish, often reflecting variation in compound type, dosing protocols, and behavioral assays [1]. In our study, ketamine induced acute, concentration-dependent behavioral alterations in adult zebrafish that were robust and reproducible. In the SPT, ketamine reduced social interaction and increased locomotor activity across all concentrations, while in the OTT it led to more selective effects, including transient increases in rotational behavior and exploratory movement. Despite repeated exposures, no behavioral sensitization emerged, and ketamine-induced changes dissipated following a 48-hour washout in both 5- and 14-day protocols. If anything, the OTT findings from Experiment I may point to a potential tolerance effect, as certain behaviors observed on day 1 diminished over subsequent experimental days. However, it remains unclear whether this reflects a true drug tolerance effect or merely results from repeated testing. These results suggest that, under the exposure conditions used here, ketamine’s effects are reversible and restricted to the acute phase. Although no persistent alterations were observed, we cannot rule out the possibility that zebrafish might display chronic or progressive phenotypes under alternative experimental designs.

Experiment I revealed a clear, concentration-dependent profile of behavioral alterations following ketamine exposure, consistent with schizophrenia-relevant phenotypes. In the SPT, zebrafish showed a robust decrease in social preference across all concentrations, indicated by reduced time and percent distance in the interaction zone. This phenotype parallels the social withdrawal modeled as a negative symptom in rodent studies and is consistent with effects of NMDA receptor antagonists in mammals [36–41]. Alongside reduced social interaction, ketamine produced hyperlocomotion, evidenced by increased distance traveled, swimming speed, and zone crossings. Increased locomotor activity following NMDA receptor antagonism has been widely used as a behavioral readout for the positive symptom dimension of schizophrenia in animal models [2, 42, 43]. These behavioral changes are in line with findings from other NMDA antagonists in zebrafish, which similarly induce both social withdrawal and hyperlocomotion [18, 19, 44–47]. Together, the results highlight the reproducibility and sensitivity of the SPT for detecting schizophrenia-relevant phenotypes in this model.

The OTT revealed a more selective profile. Rotational behavior, often interpreted as a marker of stereotypy or repetitive movement, increased transiently at higher concentrations but was not sustained beyond early sessions. Similar increases in rotational behavior have also been reported in previous studies of ketamine exposure, indicating a reproducible effect on repetitive motor activity [21, 22, 48]. Absolute turn angle remained unaffected, and ketamine’s impact on general locomotor outcomes in the OTT was milder than in the SPT, emphasizing the influence of test context. Previous findings indicate that NMDA antagonist-induced hyperlocomotion is modulated by environmental novelty, with stronger effects typically observed in unfamiliar settings or in response to novel stimuli [18, 46]. In our study, hyperlocomotion following ketamine exposure was greater in the SPT compared to the OTT, likely due to the contextual salience of the task, in which the animal is placed adjacent to a compartment containing ten conspecifics that may enhance arousal or motivational drive. Since the OTT was always conducted after the SPT, we cannot rule out the possibility that this difference reflects a testing order effect. However, prior experiments using MK-801 in which SPT and OTT were conducted in separate groups of animals rather than sequentially in the same individuals still revealed hyperlocomotion in the SPT but no change in distance traveled in the OTT, suggesting that the SPT may be inherently more sensitive to detecting locomotor alterations [18]. Similarly, studies using ketamine in the novel tank test, a context involving only exposure to a new arena, have also failed to detect hyperlocomotion, further supporting the idea that the expression of this phenotype is highly dependent on the salience of the testing environment [22, 49].

At the lowest concentration (10 mg/L), ketamine elicited an anxiolytic-like response, reflected by greater center time and distance in the OTT. Other studies using ketamine have also reported anxiolytic-like effects, but typically at higher concentrations than those tested here (20–60 mg/L), particularly in the novel tank test and light-dark test [20, 22]. However, these findings require further characterization, as ketamine-induced changes in locomotor activity may confound anxiety-related outcomes. In rodents, ketamine has also been shown to produce anxiolytic- or antidepressant-like effects [50–52], though some studies suggest this may only emerge in animals previously exposed to stress protocols [53, 54]. Whether a similar stress-dependent effect exists in zebrafish remains to be tested.

Unlike the study that inspired Experiment I [27], repeated ketamine exposure did not induce behavioral sensitization. Instead, responses remained consistent across test days, contrasting with rodent models where repeated NMDA receptor blockade frequently produces progressively amplified behavioral effects [26, 28, 55–57]. This is in line with previous zebrafish research showing no evidence of tolerance or sensitization in circling behavior following five days of ketamine exposure, suggesting a stable behavioral profile over repeated administrations [48].

Experiments II and III assessed behavioral persistence after 5-day and 14-day exposure schedules, each followed by a 48-hour washout. In both cases, zebrafish behavior returned to baseline, with no lasting effects observed in social interaction, locomotion, or repetitive activity. A small reduction in maximum speed following 5-day exposure and increase in immobility was detected following 14-day exposure, but overall, the results suggest that ketamine’s behavioral effects are transient in adult zebrafish under these conditions.

Studies examining the sustained effects of ketamine in zebrafish remain scarce. One study using the novel tank test reported that repeated exposure to ketamine (40 mg/L for seven days, with testing 24 hours later) did not alter general locomotor or anxiety-like behavior but increased the number of full-body rotations [49]. However, this test was conducted using a front-facing camera, which may limit the accurate detection of rotational behavior compared to our approach. In our study, rotations were assessed in the OTT using a top-down view, allowing for more precise measurement of body orientation. Our findings also differ from rodent models, where repeated ketamine exposure often results in sustained disruptions in behavior [25, 58].

Zebrafish show sustained neurogenesis and regenerative ability in the central nervous system from early development through adulthood [59–62]. This may have contributed to the lack of sensitization or long-term behavioral effects observed in our study, as the animals may recover from repeated acute exposures. Supporting this, other studies have also found no lasting damage when ketamine was administered during early developmental stages [63]. In rodents, repeated exposure to NMDA receptor antagonists like ketamine causes lasting impairment in cortical inhibitory circuits by triggering a redox-sensitive pathway. Repeated dosing increases IL-6 levels in the brain, which activates NADPH oxidase (*Nox2*) and leads to oxidative stress that disrupts the function of parvalbumin-expressing interneurons [23, 64]. These effects do not occur after a single exposure and appear only after repeated challenges. While reversible in adult rodents, they become permanent when the exposure occurs during development [64, 65]. Zebrafish may be less sensitive to these changes, potentially requiring more frequent or intense exposure to produce similar long-term effects.

The NMDA receptor is highly conserved across vertebrate species, including zebrafish [45, 66, 67]. Subunit composition, binding affinity, and overall receptor function are similar enough to support comparable acute effects across species. Therefore, differences in behavioral outcomes are unlikely to result from receptor pharmacology alone. One possible explanation involves species-specific differences in ketamine metabolism. Although zebrafish express cytochrome P450 enzymes involved in drug processing [68, 69], little is known about how ketamine is metabolized in this species. It is unclear how quickly the drug is metabolized, which metabolites are generated, and whether any of them contribute to behavioral effects. These factors may influence the limited duration of responses observed in this species. Investigating ketamine’s pharmacokinetics in zebrafish will be important for interpreting its behavioral impact and for evaluating the model’s relevance to other systems.

The findings presented here highlight the utility of zebrafish for modeling behavioral changes induced by NMDA receptor antagonists. However, to improve the translational relevance of this model, future studies should include pharmacological validation using clinically effective antipsychotic drugs and incorporate neurochemical or physiological analyses to clarify underlying mechanisms. Expanding the behavioral assessments to cover cognitive and affective domains, alongside systematic variation in exposure conditions such as dosing schedules, developmental timing, and environmental context, will allow for a more comprehensive evaluation of induced phenotypes.

The lack of sensitization or lasting behavioral effects likely reflects the specific exposure protocols used, rather than an inherent limitation of the zebrafish model. Variations in exposure timing, duration, dose, or the inclusion of environmental stressors may be necessary to produce more persistent effects. The zebrafish offers significant experimental flexibility, and its potential to model chronic or progressive features of psychiatric disorders warrants further investigation.

In summary, ketamine induced robust, context-dependent, and reversible behavioral alterations in adult zebrafish that parallel core symptom dimensions of schizophrenia. The use of complementary behavioral assays enabled a multidimensional assessment of drug effects, reinforcing the model’s potential for acute psychotomimetic screening. While the current exposure protocols did not produce sensitization or long-lasting behavioral changes, these findings establish a strong foundation for future studies aimed at optimizing the zebrafish model for translational neuropsychiatric research. With further refinement and validation, zebrafish hold considerable promise as a scalable platform for mechanistic discovery and early-phase therapeutic screening in drug development.

## AUTHOR CONTRIBUTIONS

**Conceptualization:** M.G.-L., D.V.M., and A.P.H. **Data curation:** M.G.-L. and A.P.H. **Formal analysis:** M.G.-L. and A.P.H. **Funding acquisition:** A.P. and A.P.H. **Investigation:** M.G.-L., D.V.M., T.S.-B., L.M.B., S.Z.B., and S.M.B. **Methodology:** M.G.-L., D.V.M., and A.P.H. **Project administration:** M.G.-L. and A.P.H. **Resources:** A.P. and A.P.H. **Supervision:** M.G.-L. and A.P.H. **Validation:** M.G.-L. and A.P.H. **Visualization:** M.G.-L. and A.P.H. **Writing - original draft:** M.G.-L. **Writing - review & editing:** D.V.M., T.S.-B., L.M.B., S.Z.B., S.M.B., A.P., and A.P.H.

## INTEREST STATEMENT

The authors state that there are no conflicts of interest related to this work.

## DATA AVAILABILITY

All data supporting this study are publicly accessible via the Open Science Framework (osf.io/67y45) [30].

## ACKNOWLEDGMENTS

The authors acknowledge the support of the Conselho Nacional de Desenvolvimento Científico e Tecnológico (CNPq, process number 305968/2023-8), the Coordenação de Aperfeiçoamento de Pessoal de Nível Superior – Brasil (CAPES), and the Pró-Reitoria de Pesquisa (PROPESQ) at the Universidade Federal do Rio Grande do Sul (UFRGS) for their funding and institutional support.

## Notes

https://doi.org/10.17605/OSF.IO/67Y45

### Competing Interest Statement

The authors have declared no competing interest.

