## Supplementary material for "Ketamine-induced NMDA receptor hypofunction alters social and locomotor behavior in adult zebrafish"

**Supplementary Figure 1**

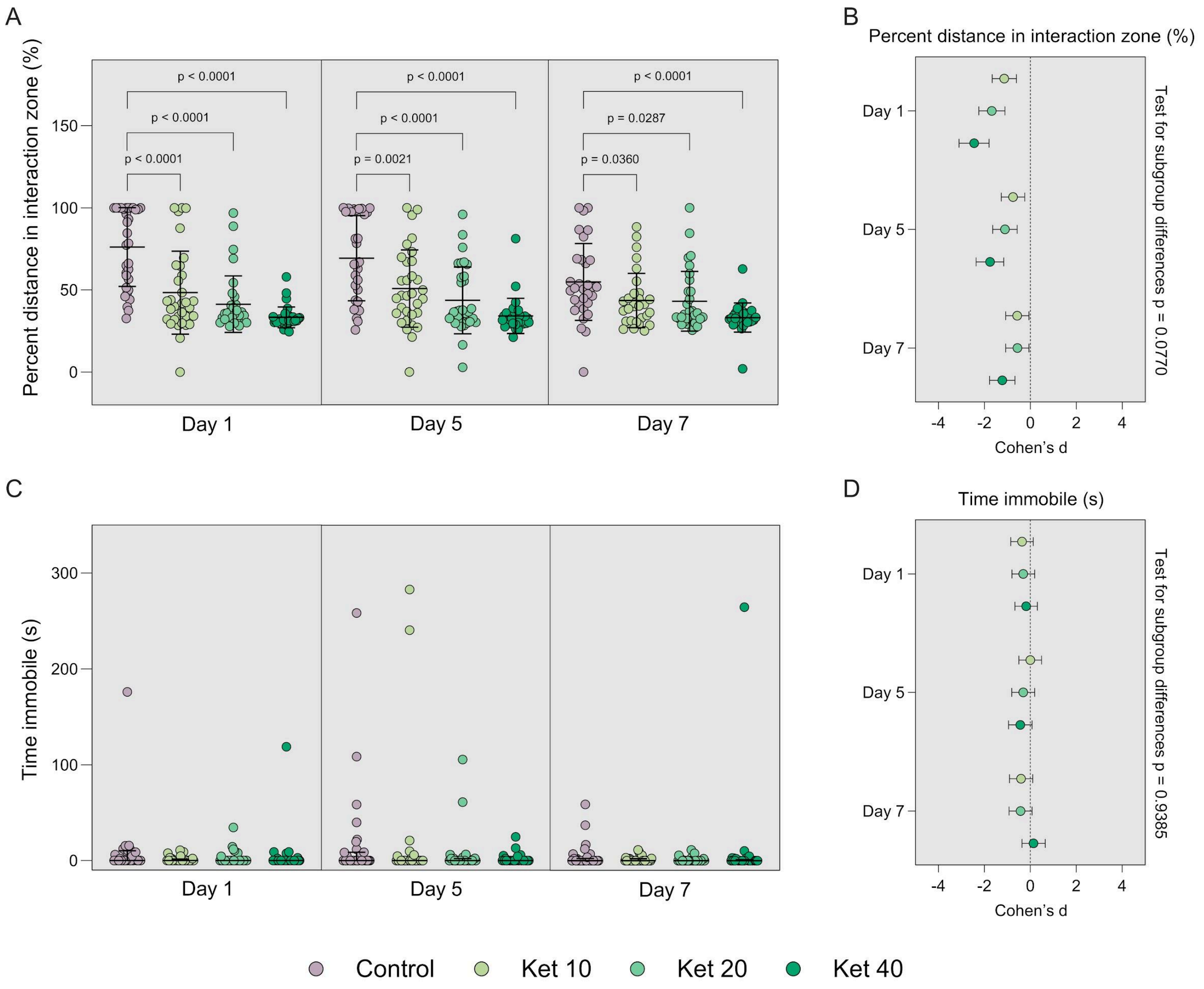


**Supplementary Figure 1.** Additional behavioral outcomes from the social preference test in adult zebrafish exposed to different concentrations of ketamine. (A) Percent of distance traveled in the interaction zone and (C) time spent immobile, measured on days 1, 5, and 7. (B, D) Corresponding effect sizes (Cohen's d) for each outcome, with test for subgroup differences across assessment days. One-way ANOVA followed by Bonferroni post hoc test was used for A; Kruskal–Wallis test followed by Dunn's post hoc was used for C. Data are presented as mean ± SD for A, and as median with interquartile range for C. Effect sizes are expressed as Cohen's d with 95% confidence intervals. Subgroup differences tested using a random-effects model. Sample sizes: day 1 – Control = 32, Ket 10 = 32, Ket 20 = 32, Ket 40 = 32; day 5 – Control = 31, Ket 10 = 32, Ket 20 = 32, Ket 40 = 29; day 7 – Control = 31, Ket 10 = 31, Ket 20 = 31, Ket 40 = 29. Ket 10 = ketamine 10 mg/L; Ket 20 = ketamine 20 mg/L; Ket 40 = ketamine 40 mg/L.

### Supplementary Figure 2

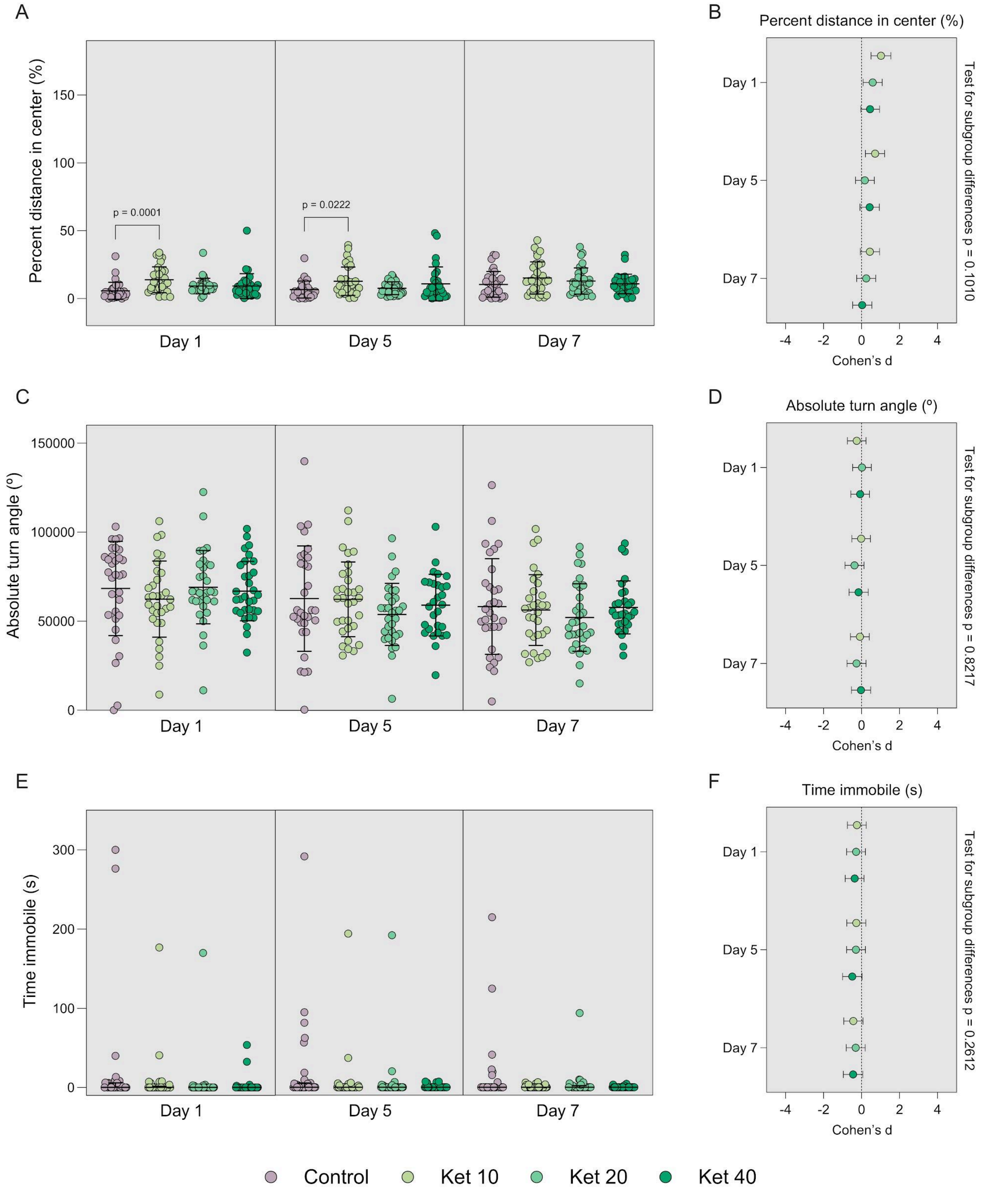


**Supplementary Figure 2.** Additional behavioral outcomes from the open tank test in adult zebrafish exposed to different concentrations of ketamine. (A) Percent of distance traveled in the center zone, (C) absolute turn angle, and (E) time spent immobile, measured on days 1, 5, and 7. (B, D, F) Corresponding effect sizes (Cohen's d) for each outcome, with test for subgroup differences across assessment days. One-way ANOVA followed by Bonferroni post hoc test was used for A and C; Kruskal–Wallis test followed by Dunn's post hoc was used for E. Data are presented as mean ± SD for A and C, and as median with interquartile range for E. Effect sizes are expressed as Cohen's d with 95% confidence intervals. Subgroup differences tested using a random-effects model. Sample sizes: day 1 – Control = 32, Ket 10 = 32, Ket 20 = 32, Ket 40 = 32; day 5 – Control = 31, Ket 10 = 32, Ket 20 = 32, Ket 40 = 29; day 7 – Control = 31, Ket 10 = 31, Ket 20 = 31, Ket 40 = 29. Ket 10 = ketamine 10 mg/L; Ket 20 = ketamine 20 mg/L; Ket 40 = ketamine 40 mg/L.
